# Group ICA for Identifying Biomarkers in Schizophrenia: ‘Adaptive’ Networks via Spatially Constrained ICA Show More Sensitivity to Group Differences than Spatio-temporal Regression

**DOI:** 10.1101/429837

**Authors:** Mustafa S Salman, Yuhui Du, Dongdong Lin, Zening Fu, Eswar Damaraju, Jing Sui, Jiayu Chen, Qingbao Yu, Andrew Mayer, Stefan Posse, Daniel Mathalon, Judith M. Ford, Theodorus Van Erp, Vince D Calhoun

## Abstract

Brain functional networks identified from fMRI data can provide potential biomarkers for brain disorders. Group independent component analysis (GICA) is popular for extracting brain functional networks from multiple subjects. In GICA, different strategies exist for reconstructing subject-specific networks from the group-level networks. However, it is unknown whether these strategies have different sensitivities to group differences and abilities in distinguishing patients. Among GICA, spatio-temporal regression (STR) and spatially constrained ICA approaches such as group information guided ICA (GIG-ICA) can be used to propagate components (indicating networks) to a new subject that is not included in the original subjects. In this study, based on the same a priori network maps, we reconstructed subject-specific networks using these two methods separately from resting-state fMRI data of 151 schizophrenia patients (SZs) and 163 healthy controls (HCs). We investigated group differences in the estimated functional networks and the functional network connectivity (FNC) obtained by each method. The networks were also used as features in a cross-validated support vector machine (SVM) for classifying SZs and HCs. We selected features using different strategies to provide a comprehensive comparison between the two methods. GIG-ICA generally showed greater sensitivity in statistical analysis and better classification performance (accuracy 76.45±8.9%, sensitivity 0.74±0.11, specificity 0.79±0.11) than STR (accuracy 67.45±8.13%, sensitivity 0.65±0.11, specificity 0.71±0.11). Importantly, results were also consistent when applied to an independent dataset including 82 HCs and 82 SZs. Our work suggests that the functional networks estimated by GIG-ICA are more sensitive to group differences, and GIG-ICA is promising for identifying image-derived biomarkers of brain disease.

## 1 Introduction

Functional brain networks derived from functional magnetic resonance imaging (fMRI) data may serve as potential biomarkers for many mental disorders. One of the most widely applied multivariate methods for estimating brain functional networks is independent component analysis (ICA). Spatial ICA (V. D. Calhoun et al., 2001; Mckeown et al., 1998), which models the fMRI data as a combination of spatially independent sources with each being related to a time course (TC), has been widely applied in functional MRI studies. Unlike traditional techniques such as general lineal model (GLM) and region of interest based methods, ICA requires no prior information in determining the regions and time series. Another advantage of ICA is that it can analyze multiple brain networks at the same time by considering the whole fMRI data, while traditional methods need to set separate prior information for extracting each of the multiple networks under experiment. ICA can also denoise the fMRI data by decomposing artifacts as independent components, thereby extracting more meaningful components. However, the biggest challenge in ICA comes from the arbitrary order of the obtained components. This limitation makes the functional networks of different subjects, computed by performing individual ICA on each subject’s fMRI data, not directly corresponding across subjects. Since network correspondence is necessary for statistical analysis and classification in multi-subject studies, estimating accurate subject-specific networks with the same biological meaning across subjects is a key.

Group ICA (GICA) has been proposed to solve the problem of establishing subject correspondence in multi-subject studies. It involves performing ICA on the group data by temporal concatenation (Beckmann et al., 2009; V.D. Calhoun et al., 2001), spatial concatenation (Svensén et al., 2002) or tensor organization (Beckmann and Smith, 2005; Lee et al., 2008). The temporal concatenation is by far the most widely used approach and allows for unique TCs for each subject and while assuming spatial stationarity of the maps, still does allow for considerable variability in the single subject maps (Allen et al., 2012). However, the degree to which the trade-off between a group model and an individual subject representation is traversed depends on the specific algorithm being used (Erhardt et al., 2011; Michael et al., 2014). In fMRI studies, the temporal concatenation method is widely applied (Allen et al., 2011; Calhoun and Adali, 2012; Schmithorst and Holland, 2004; Smith et al., 2013). Typical temporal concatenation-based group ICA approaches perform principal component analysis (PCA) dimension reduction on fMRI data followed by ICA, which generates group-level independent components. Next, a back-reconstruction step is implemented to estimate subject-specific components and their associated subject-specific TCs. The widely-used Group ICA of fMRI toolbox (GIFT) (Calhoun, 2004) incorporates several methods for the back-reconstruction step, including three PCA-based approaches (GICA1, GICA2 and GICA3) (Calhoun, 2004), a least-squares based approach called spatio-temporal regression (STR) or dual regression (Beckmann et al., 2009; Erhardt et al., 2011), and two spatially constrained approaches (Lin et al., 2010) including group information guided ICA (GIG-ICA) which incorporates a multiple-objective function optimization approach (Du and Fan, 2013). Among the strategies implemented in the GIFT toolbox, both STR and GIG-ICA can be used to estimate the subject-specific networks for additional subjects who are not used for computation of the group-level components, while PCA-based methods cannot be as easily extended this way. GIG-ICA simultaneously optimizes the independence among individual networks of each subject and the dependence of networks across subjects, providing a nice balance between the group model (matching of components) and the individual subject specificity of the estimated networks, including well estimated resting networks in the context of highly subject specific artifacts (Du et al., 2015b, 2016a, 2017b; Du and Fan, 2013). Spatial networks derived from ICA analysis of fMRI data are extensively used as features in classification of mental illness such as schizophrenia (Castro et al., 2011; Du et al., 2012). For classification (or diagnosis) of new subjects, it is necessary to extract accurate individual networks while still preserving network correspondence with previous subjects.

Schizophrenia is a chronic illness associated with widespread changes in brain connectivity. Meda et al. reported abnormal resting-state functional network connectivity (FNC) in schizophrenia and psychotic bipolar patients (Meda et al., 2012). Temporally coherent brain networks such as temporal lobe and default mode networks have been shown to reliably discriminate subjects with bipolar disorder, chronic schizophrenia and healthy controls (Calhoun et al., 2008). The interactions among brain networks have also been implicated in healthy population and various clinical groups. It has been shown that patients with schizophrenia tend to linger in a state of weak connectivity at rest (Damaraju et al., 2014a; Du et al., 2016b). Similar findings have been reported in patients with bipolar disorder (Rashid et al., 2014). Group ICA methods have also identified potential biomarkers for schizophrenia, bipolar disorder and schizoaffective disorder (Du et al., 2016b, 2015b, 2015b, 2015a, 2014). While Erhardt et al. (Erhardt et al., 2011) performed a comparison of back-reconstruction approaches in an extensive simulation study, no studies have directly compared back-reconstruction approaches in the context of biomarker detection from real data. Thus, it is unclear which back-reconstruction method is more sensitive in revealing subtle differences between groups or subjects when the goal is to translate the results to new datasets (e.g. for an identified set of biomarkers).

In this paper, we compare the STR and GIG-ICA methods in terms of their ability to identify brain network-based biomarkers that can discriminate healthy controls (HCs) from schizophrenia patients (SZs). Our hypothesis was that GIG-ICA would be more sensitive, as the use of STR on new subjects assumes fixed component maps (Joel et al., 2011), whereas GIG-ICA re-optimizes the component maps given the new data, while also preserving the component ordering. In our study, we first estimated the group-level components from the publicly available FBRIN dataset using ICA. Using these group-level networks as prior, we used both STR and GIG-ICA methods to back-reconstruct the subject-specific functional networks from the fMRI data of the FBIRN dataset and then investigated the group differences on the spatial networks and the FNC for each method. We also applied the support vector machine (SVM) machine learning technique to classify HCs and SZs in the FBIRN dataset using the networks as features. To corroborate this classification outcome, we used the same priors to estimate subject-specific networks from the COBRE dataset consisting of HC and SZ samples independent of the FBIRN dataset, and then performed classification on it.

## 2 Materials and Methods

### 2.1 Materials

We employed data from the Functional Biomedical Informatics Research Network (FBIRN) phase-III study. Resting-state fMRI data were originally collected from 186 healthy controls (HCs) and 176 schizophrenia patients (SZs). The subjects were age and gender-matched. The SZ subjects were diagnosed using Structured Clinical Interview for DSM-IV-TR Axis I Disorders (First et al., 2002). The exclusion criteria for the SZ subjects was based on a current or past history of major medical illness and having significant extrapyramidal symptoms or tardive dyskinesia or significant changes in psychotropic medications in the previous two months before the scan. Any healthy subject with a current or past history of major neurological or psychiatric medical illness, or a first degree relative with a psychotic illness diagnosis was also excluded. Subjects were also excluded if they did not have any of the following: normal hearing levels, sufficient eyesight to see visual displays, IQ greater than 75, fluency in English and ability to perform the study tasks, or if they had previous head injury or prolonged unconsciousness, substance or alcohol dependence, migraine treatments or MRI contradictions. In the resting-state fMRI scan, 162 volumes of echo planar imaging (EPI) BOLD fMRI data were collected on 3T scanners in eyes closed condition with the following imaging parameters: FOV = 220mm×220mm (64×64 matrix), TR = 2s, TE = 30ms, flip angle = 77^0^, 32 sequential ascending axial slices with thickness of 4mm and 1mm skip (Keator et al., 2016).

The independent dataset used in external classification evaluation came from the Center of Biomedical Research Excellence (COBRE) study conducted at the Mind Research Network (MRN). Resting-state fMRI data of 100 HCs and 87 SZs consisted of 149 volumes of T2*-weighted functional images each, acquired using a gradient-echo EPI sequence: TR = 2 s, TE = 29 ms, flip angle = 75°, slice thickness = 3.5 mm, slice gap = 1.05 mm, field of view = 240 mm, matrix size = 64 × 64 and voxel size = 3.75 mm × 3.75 mm × 4.55 mm.

### 2.2 Methods

#### 2.2.1 Data Preprocessing

FMRI data of FBIRN were preprocessed using the SPM (Friston, 2007) and AFNI (Cox, 1996) toolboxes. The initial 6 volumes from each scan were discarded to allow for equilibration of T1-related signal saturation. Next, the signal-fluctuation-to-noise ratio (SFNR) of all subjects was calculated (Friedman et al., 2006). In addition, the INRIAlign (Freire et al., 2002) toolbox in SPM was used to perform rigid body motion correction, which produced a measure of maximum root mean square (RMS) translation. All subjects with SFNR<150 and RMS translation>4mm were excluded (Damaraju et al., 2014a). The remaining 314 subjects, consisting of 163 healthy controls (HCs; mean age 36.9, 46 females) and 151 schizophrenia patients (SZs; mean age 37.8, 37 females) were included for further analysis. Next, slice-timing correction was performed to account for timing difference in slice acquisition (Damaraju et al., 2014a). The fMRI data were then despiked using AFNI’s 3dDespike algorithm to reduce the effect of outliers (Damaraju et al., 2014b). The images were subsequently spatially normalized to the standard Montreal Neurological Institute (MNI) space and resampled to 3mm×3mm×3mm voxels. Finally, the images were smoothed to 6mm full width at half maximum (FWHM). Additional details about data preprocessing can be found in (Damaraju et al., 2014a). The same preprocessing procedure as above was used while preprocessing the fMRI data of COBRE. After preprocessing, a total of 164 subjects (82 HCs, 82 SZs) out of 187 were retained for further analysis.

#### 2.2.2 Group-level component computation

In this work, we used the 47 group-level network-related components from our previous study (Damaraju et al., 2014a) as prior, obtained from the data of FBIRN. These group-level networks were estimated by performing group-level ICA on the temporal concatenation of preprocessed fMRI data of all subjects (V.D. Calhoun et al., 2001). The procedure is presented in Fig. 1(A) which includes two steps: (1) performing the subject-level principal component analysis (PCA) with the number of principal components as 120 (PC1 = 120) on each subject’s fMRI data and group-level PCA with the number of principal components as 100 (PC2 = 100) on the reduced and concatenated data, and (2) performing ICA with Infomax (Bell and Sejnowski, 1995) on the PCA-reduced data, resulting in group-level components. To ensure the stability of IC estimation, ICA was repeated 20 times in ICASSO (Ma et al., 2011) and the 100 most reliable components were identified as the final group-level components. Finally, the 100 independent components were evaluated to identify the resting-state networks. The criteria for identifying the networks were: a. the peak activation clusters of a network should be in grey matter, b. there should be minimal overlap with known vascular, susceptibility, ventricular and edge regions, and c. the mean power spectra of the networks should show higher low frequency spectral power. Following this selection procedure, 47 resting-state networks were obtained out of 100 spatially independent components.

#### 2.2.3 Reconstruction of individual networks using STR and GIG-ICA

For the FBIRN data, the back-reconstruction step involves estimating subject-specific networks and their associated time courses (TCs) for each of the 314 subjects based on the selected 47 group-level networks. In this study, we performed back-reconstruction using spatio-temporal regression (STR) and group information guided ICA (GIG-ICA) separately. Both methods can be implemented using the GIFT toolbox (Calhoun, 2004). Note that the subject-specific networks reconstructed by both methods were based on the same prior networks, and thereby were corresponding across individual networks as well as comparable across the two methods.

STR uses a least squares approach to estimate subject-specific networks and their associated TCs (Beckmann et al., 2009; Erhardt et al., 2011). This approach is as follows. Let **Y**_***i***_ = **R**_***i***_**S** + **E**_1***i***_, where **Y**_***i***_ is the *T* × *V* matrix for subject *i* (*i* = 1,2, …, *M*) after preprocessing, *T* indicates time points and *V* is the number of voxels, **R**_***i***_ is the subject-specific TC, **S** is the matrix of estimated group-level networks and **E**_1***i***_ indicates the error. In the first step, as shown in Fig. 1(C1), least squares estimation for the TC 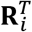 gives 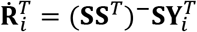. Next, the subject-specific networks, **S**_***i***_ are estimated from the TCs 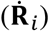. Here, the earlier assumption that all subjects share a common network, is relaxed (Erhardt et al., 2011). According to Fig. 1(C2), let 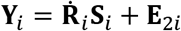. Then least squares estimation for **S**_***i***_ gives the subject networks: 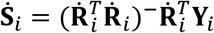. After obtaining the individual **S**_***i***_, each network (row of **S**_***i***_) was z-scored to zero mean and unit variance.

We compare STR and GIG-ICA methods in this work because they allow estimation of new subject-specific networks (and TCs) from a set of reference ICs. Hence, the **S** matrix above consists of 47 columns of selected group-level resting state networks out of 100 ICs. However, it has been argued that the inclusion of artifactual components improves the STR estimation. Therefore, we also estimated the subject-specific networks by including all 100 ICs in the STR estimation. We then selected the 47 subject-specific networks corresponding to the group-level resting state networks in the subsequent experiments and reported the classification results corresponding to those features separately in the results section.

GIG-ICA using multi-objective optimization framework is proposed to estimate individual networks (Du et al., 2015b; Du and Fan, 2013). Let **S**^*k*^ denote the *k*^*th*^ group-level reference IC. The optimization, as depicted in Fig. 1(B), is 

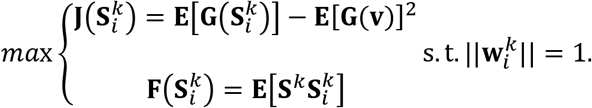

Here, 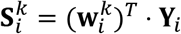 denotes the *k*^*th*^ subject-specific IC of the *i*^*th*^ subject, which corresponds to **S**^*k*^. **Y**_***i***_ denotes the random vector of whitened fMRI data of the ***i***^*th*^ subject, and 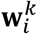 is the unmixing vector. 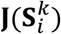, the negentropy of the estimated 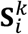 with updates on 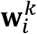, serves to measure the independence of the IC. **G**(⋅) is a nonquadratic function and **v** is a Gaussian variable with zero mean and unit variance. 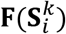 measures the similarity between **S**^*k*^ and 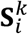, with **E**[] denoting the expectation of variable. Solving the optimization function results in the optimal 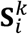. The algorithm automatically generates z-scored 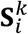, which can be compared across subjects. Subsequently, the subject-specific TC can be computed as 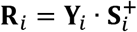, where 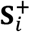 denotes the pseudo-inverse matrix of the estimated components.

**Fig. 1:**
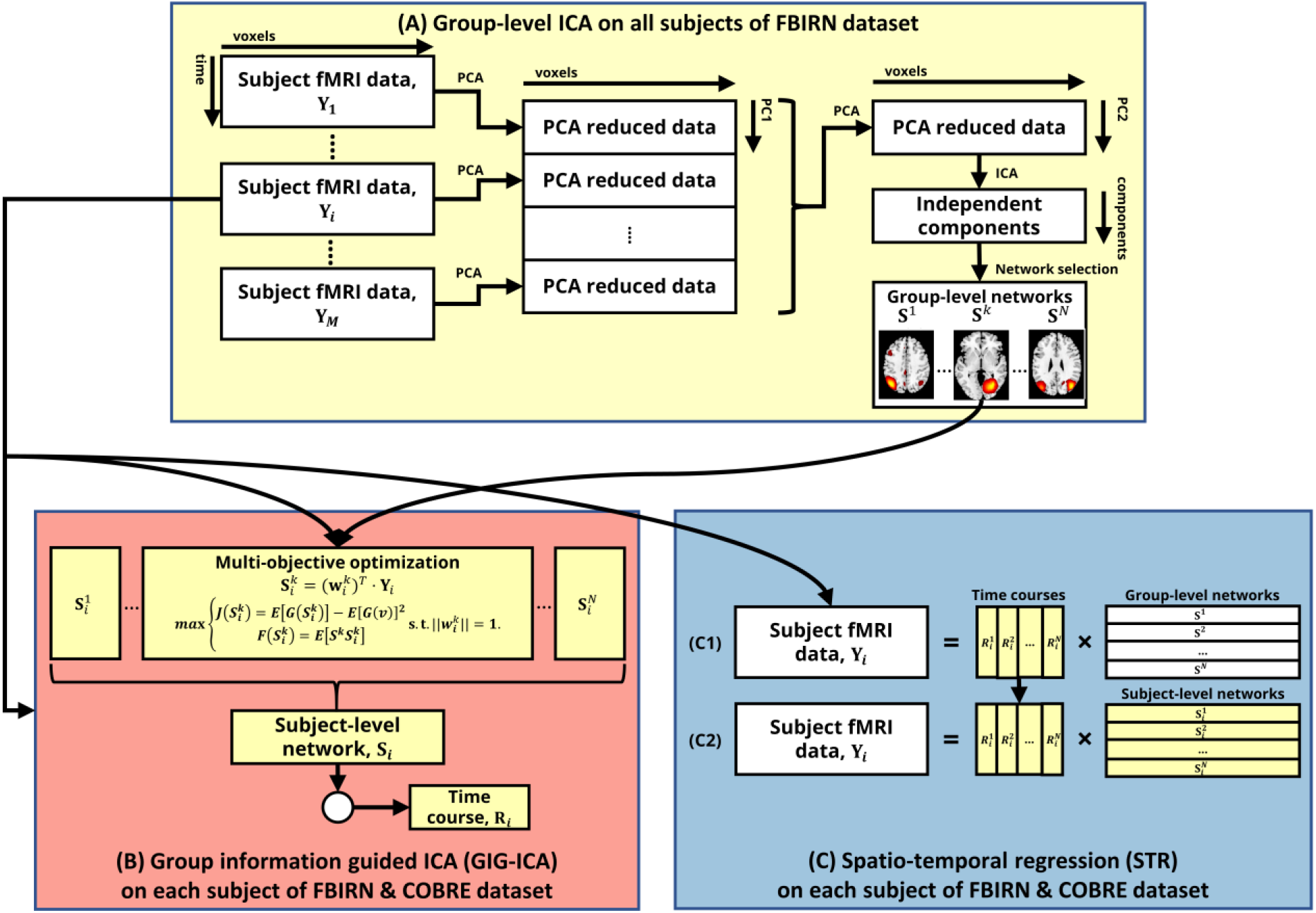
Method framework of (A) group-level ICA on all subjects’ fMRI data to estimate the group-level networks, (B) group information guided ICA (GIG-ICA) and (C) spatio-temporal regression (STR). (B) and (C) are performed on each subject’s fMRI data to estimate the subject-specific networks and time courses.

#### 2.2.4 Comparison between STR and GIG-ICA for distinguishing HCs and SZs

The subject-specific networks and TCs estimated using a back-reconstruction method such as GIG-ICA or STR allows us to perform group comparison by statistical analyses. The following sections describe the analyses undertaken to compare the HC vs. SZ group differences in the functional networks and the functional network connectivity (FNC) for each method. A machine learning method is also applied to classify HCs and SZs using the spatial networks estimated by each method, aiming to compare which back-reconstruction strategy can better differentiate patients and controls at the level of the single-subject.

##### 2.2.4.1 Comparing group difference in the functional network maps

Different strategies were used to provide a comprehensive comparison of the methods. Fig. 2 presents three different strategies for testing group difference between the subject networks. For strategy 1, firstly within each network estimated by using GIG-ICA and STR, a right-tailed one sample t-test was performed on the z-score of each voxel across all subjects. The voxels with significantly positive z-score (*p* < 0.01 after Bonferroni correction for multiple comparison) in both methods were identified, and then an overlap of the significant voxels in both methods was chosen. This resulted in one common mask for each network. Next, on every voxel z-score within this mask, a two-sample t-test was performed to test the group difference between HC and SZ. The voxels with significant group differences (*p* < 0.05, uncorrected) in both networks estimated by GIG-ICA and STR were identified by another overlap in strategy 1. The p-value threshold was chosen to retain at least one voxel within every network for subsequent comparison between the methods. Note that for a particular network, some voxels show significantly higher z-scores in HC than SZ, whereas for the other voxels within the same network it can be the opposite. The first set of voxels is referred to as *V*_*HC*>*SZ*_, whereas the latter is referred to as *V*_*HC*<*SZ*_. Strategy 2 differed from the previous one in that no overlap of significant voxels was taken after either the one sample t-test step or the two-sample t-test step. Therefore, the processes of significant voxel identification for GIG-ICA and STR were independent between the methods. Whereas in strategy 3, an overlap of significant voxels with positive z-scores was taken after the one sample t-test step, while the two-sample t-test step (group comparison) was performed for each method independently.

In the three strategies discussed above, within each network, the discriminatory voxel z-scores estimated by the two back-reconstruction methods were compared separately. Afterwards, using a permutation test, the significant difference between the voxel t-stats estimated by GIG-ICA and STR were obtained at the network level. Within a network, the t-stats obtained using GIG-ICA were assigned factor=1 and the ones from STR were assigned factor=0. The difference of mean of t-stats with factor=1 and mean of t-stats with factor=0 was noted. The factors were then permuted 10000 times and a distribution of the observed mean difference was built. This distribution was used to obtain a p-value for the mean difference via two-tailed test. The statistical difference between t-stats obtained from GIG-ICA and STR in each network was noted.

**Fig. 2:**
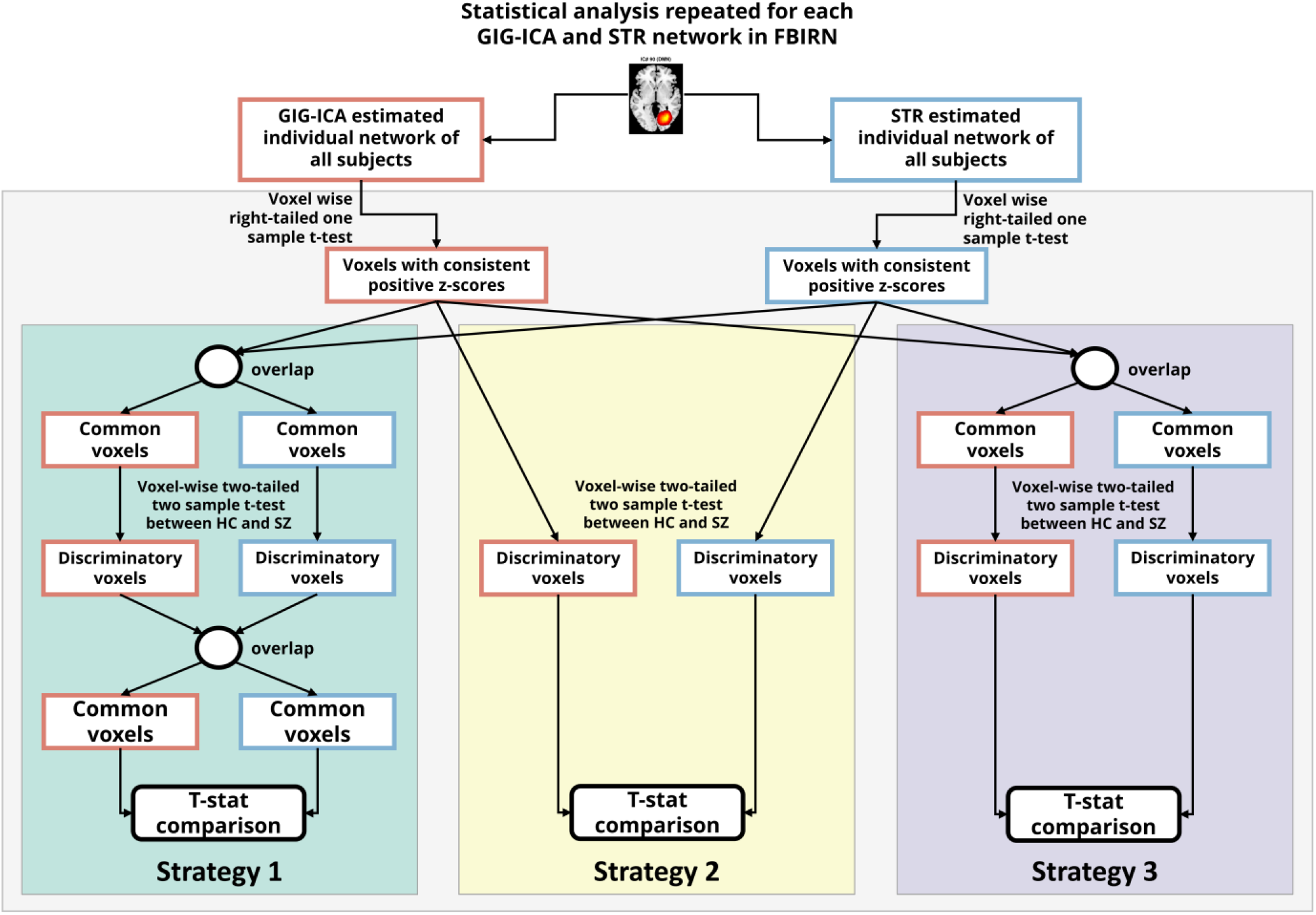
Statistical analysis procedure on brain functional networks. For each network estimated by GIG-ICA and STR, three different strategies were employed to identify the voxels showing significant effect of diagnosis. In strategy 1, after both one-sample t-test and two-sample t-test, the common significant voxels between GIG-ICA and STR were considered. In strategy 2, common significant voxels between the two methods were not considered at any step (hence the comparison was independent between the methods). In strategy 3, the common significant voxels between the methods were considered after one-sample t-test, but not after two-sample t-test.

##### 2.2.4.2 Classification based on the functional network maps

For FBIRN data, classification was performed by using a 10-fold cross-validated training-testing framework with 100 repeats. In each repeat, randomly selected 90% of FBRIN data (*N_train_*) were used for selecting features and training a classifier model, and then individual-subject classification differentiating controls and patients was implemented for the remaining testing data (*N_test_*). The framework is described in Fig. 3(A). Three different strategies (as described in sec. 2.2.4.1) allowed extracting three different sets of discriminatory voxels from each method for the training data. The corresponding z-scores (in the subject-specific ICs) were used as the features to train an SVM classifier with radial basis function (RBF) kernel, which was then used to predict the testing set of subjects (HC vs. SZ). As SVM is not scale-invariant, each feature of the training set and testing set were standardized across subjects separately. The classification method is detailed below.

A C-support vector classifier (C-SVC, C is the regularization parameter of the SVM algorithm) with RBF kernel was used to classify the controls and the patients using the features identified in the previous step. The LibSVM toolbox for MATLAB was used to perform the classification (Chang and Lin, 2011). The classification experiment was repeated 100 times for GIG-ICA and STR estimated network features and each of the three strategies to obtain a stable measure of the classification results, each time with randomly selected training/testing samples. In each of these repeats, a 10-fold cross-validation was performed within the training subset (*N_train_*) to determine the optimum cost parameter (*C*) and kernel parameter (*γ*) of the classifier as follows. The training samples were divided into approximately 10 equal folds. In each iteration (out of 10), one of the folds was set aside for testing and the features from the other 9 folds were used for model training. As a result, every subject in the original training set was used at least once for testing the model during cross validation. Initially a coarse cross-validation was performed, i.e. the *C* parameter was selected from 10 logarithmically spaced values between 10 to 10^15^ and *γ* from another 10 values between 10^−15^ to 1. The values were logarithmically spaced to reduce the amount of time required to run cross-validation. Once an optimum pair of parameters (corresponding to the highest testing accuracy, the ratio of correctly labelled subjects and total number of subjects in the testing set) was determined from the logarithmically spaced sequence, a fine cross-validation was performed based on linearly spaced parameter values in the proximity of the identified optimum parameter. It was possible to obtain the optimum parameter with reasonable cross-validation accuracy from these steps. For each of the 10 folds, an SVM model was generated using the optimum pair of (*C, γ*) parameters and the features extracted from the training set. The model was then used for predicting the test accuracy from the remaining fold. The (*C, γ*) pair for which the validation accuracy was highest was retained. Finally, the selected features and the (*C, γ*) parameters from the training set were used to predict the test accuracy from the remaining *N_test_* subjects.

**Fig. 3:**
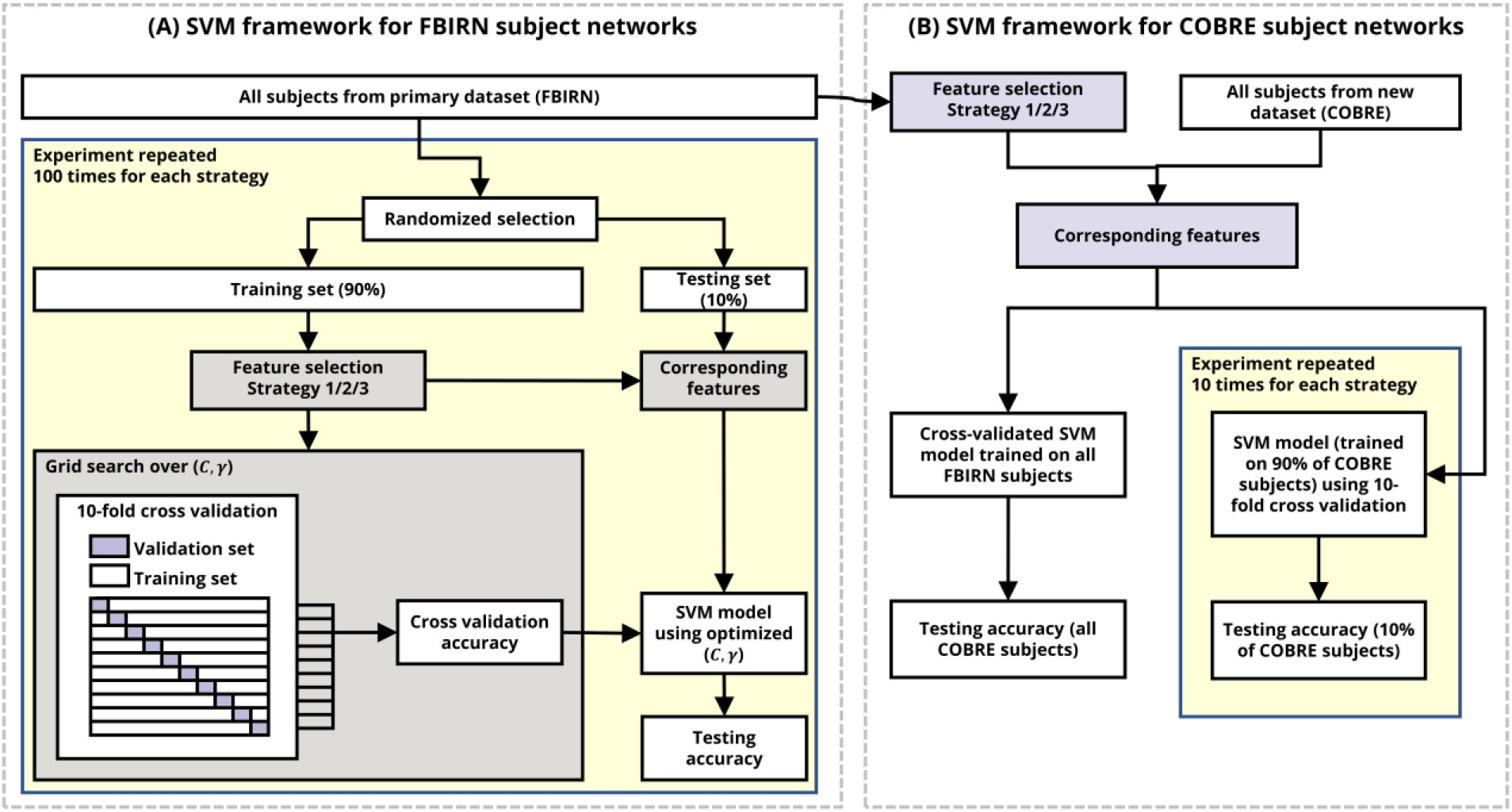
Framework for SVM classification on GIG-ICA and STR estimated networks. (A) Classification framework using the (primary) FBIRN data with 100 times of 10-fold cross-validation. In each time, a random training set was selected, on which 3 strategies were used for feature selection. Three sets of selected features were used separately to train SVM models whose optimum parameters (C,\gamma) were determined by another 10-fold cross-validation within the training data. Then the model was used to predict the testing set using the same features. (B) Classification framework using the COBRE data, independent from the primary FBIRN data. Two different schemes were applied to evaluate the classification ability in COBRE data. In the first scheme, an SVM model was trained using all the subjects from the FBIRN dataset, and this model was used to predict the labels for each of the subjects in the COBRE dataset. In the second scheme, 10-fold cross-validation within COBRE dataset was performed.

The generalizability of the identified biomarkers employed to distinguish SZs was also examined and compared between GIG-ICA and STR using the independent COBRE dataset. The framework is described in Fig. 3(B). For each subject in COBRE data, the subject-level networks were computed (using GIG-ICA and STR) based on the group-level networks resulting from section 2.2.2 and its preprocessed fMRI data. The estimated networks of subjects in FBIRN and COBRE had the same order, therefore the features can be correspondingly extracted. It is worth noting that the feature selection was performed only based on the FBIRN data, and three strategies were applied in this step. For COBRE data, two different classification schemes were employed. In the first scheme, an SVM model was trained using all the subjects from the FBIRN dataset, and this model was used to predict the labels for each of the subjects in the COBRE dataset. In the second scheme, an SVM model was trained using 90% of the COBRE dataset, which was then used to predict the rest of the subjects in COBRE, and this procedure was repeated 10 times with different sets of randomly selected training/testing subjects. In both schemes, the optimal parameters of SVM were estimated using 10-fold cross-validation within the training data.

##### 2.2.4.3 Comparing the functional network connectivity

FNC assesses between-network connectivity, i.e. the interaction among networks. In the ICA approach, this can be studied by examining the temporal relationship among the associated TCs of the networks. In our work, the group difference in FNC was investigated using FBIRN data as follows. The subject-specific TCs of the functional networks were first post-processed by removing linear, quadratic and cubic trends, regressing out head motion parameters and despiking using AFNI’s 3dDespike algorithm (Cox, 1996), which reduced the impact of outliers on FNC computation. Correlation among brain networks has been shown to be primarily driven by low frequency fluctuations in BOLD fMRI data (Cordes et al., 2001). Hence the network TCs were also filtered with a 5^th^ order low-pass Butterworth filter with a high cut-off frequency of 0.15Hz. To reflect the interaction among networks, a 47 × 47 FNC matrix of each subject was computed using the Pearson correlations between the paired post-processed TCs.

A paired t-test was performed between the subject-specific connectivity estimated by GIG-ICA and STR to illustrate the contrast between the two methods. Finally, for each method, group differences between HCs and SZs were identified by performing a two-tailed two-sample t-test on each element (representing connection between two networks) in the FNC matrix. Then the identified group differences were compared between the two methods.

## 3 Results

### 3.1 Comparison results of the network maps

The 47 group-level networks are presented in Fig. 4. Based on their known anatomical and functional properties, these networks are grouped into the following functional domains: 5 subcortical (SC), 2 auditory (AUD), 10 visual (VIS), 6 sensorimotor (SM), 9 attention (ATTN), 7 frontal (FRN), 6 default-mode (DMN), and 2 cerebellar (CB) networks. More details including the network labels and coordinates of peak activation of these networks can be found in previous work (Damaraju et al., 2014a).

**Fig. 4:**
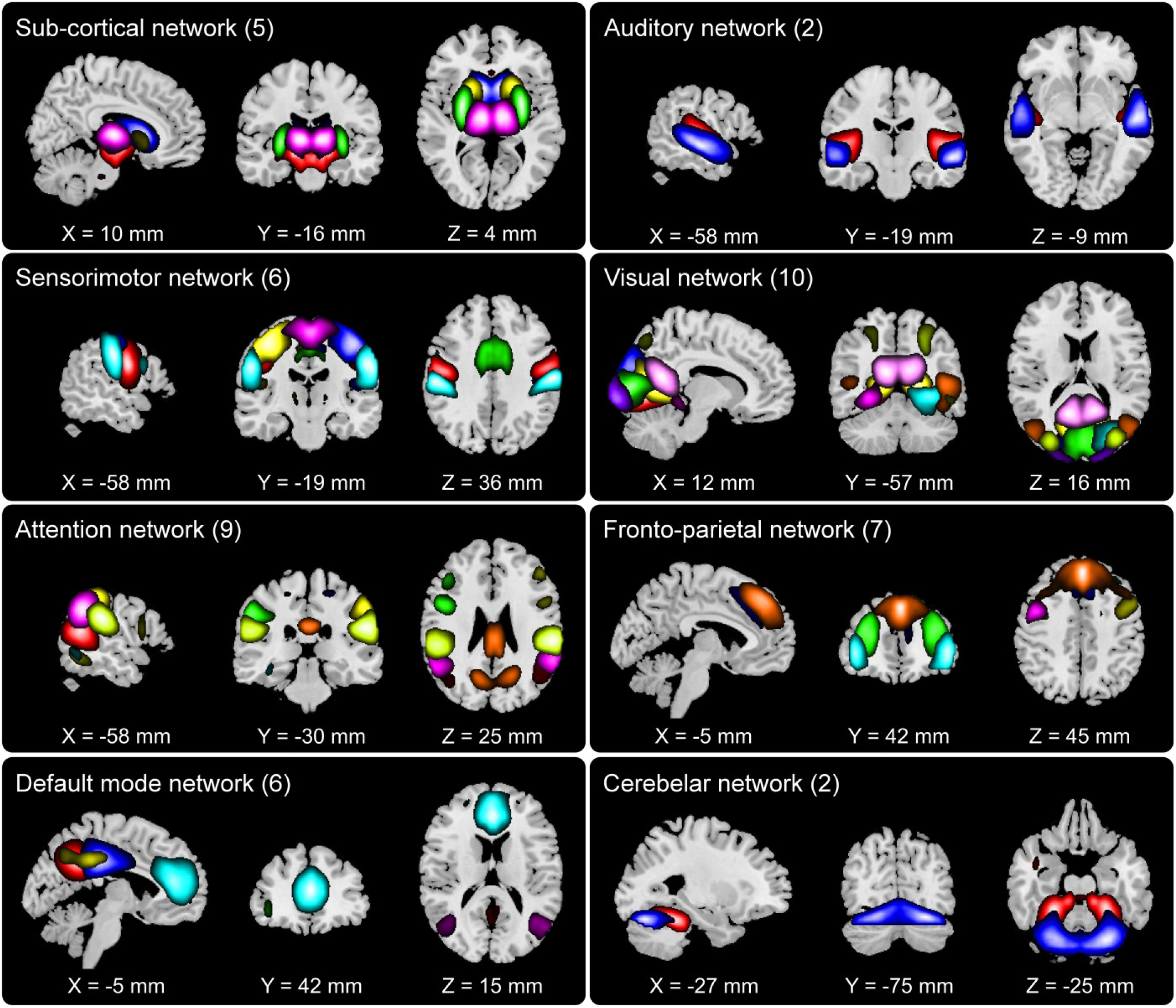
Composite view of 47 group-level networks grouped into functional domains: 5 subcortical (SC) networks, 2 auditory (AUD) networks, 10 visual (VIS) networks, 6 sensorimotor (SM) networks, 9 attention (ATT) networks, 7 fronto-parietal (FRN) networks, 6 default mode networks (DMN) and 2 cerebellar (CB) networks. Intensity of color represents z-scores. Component labels and peak activation coordinates can be found in previous work (E. Damaraju et al., 2014).

Fig. 5 displays the scatterplots of the t-stats (i.e., t-values from two-sample t-tests) within each of the 47 networks estimated using GIG-ICA and STR respectively. Three subplots show t-stats for three different sets of voxels obtained from three different strategies, as outlined in section 2.2.4.1 as well as Fig. 2. Each scatterplot has a break on the y-axis, with the top half presenting t-stats from the voxels in the *V*_*HC*>*SZ*_ mask, and the bottom half showing the results from the *V*_*HC*<*SZ*_ mask. Note that the two-sample t-test was performed by comparing HC to SZ, hence the t-stats are negative in the bottom halves of the plots (*V*_*HC*<*SZ*_ mask). Supplementary Fig. S1, S2 and S3 present the t-map of these discriminatory voxels at the slices with highest variation in t-stats, obtained from the three different strategies respectively. The individual voxel based group differences are consolidated for each network through a permutation test, the result of which are indicated by the red stars and blue circles in Fig. 5.

Fig. 5(A) shows that in strategy 1, for 31 out of 47 networks’ *V*_*HC*>*SZ*_ mask, the t-stats are significantly more positive when the networks are estimated by GIG-ICA compared to STR, as indicated by red stars on the top half of the subfigure. For one out of the other 16, namely left angular gyrus (DMN), STR shows significantly higher (more positive) t-stats than GIG-ICA, as indicated by the blue circle. For the rest of 15 networks, the t-stats were not significantly different between GIG-ICA and STR in the permutation test. For 17 networks of the *V*_*HC*<*SZ*_ mask, the t-stats are significantly more negative when estimated by GIG-ICA than STR. However, there are 6 other networks in the *V*_*HC*<*SZ*_ mask where STR shows significantly more negative mean t-stats than GIG-ICA. Fig. 5(B) shows that in strategy 2, for 34 out of 47 networks’ *V*_*HC*>*SZ*_ mask, the t-stats are significantly more positive when the networks are estimated by GIG-ICA. In contrast, there are 3 networks, namely precuneus, left angular gyrus and another angular gyrus component where the t-stats are significantly more positive when estimated by STR. In case of the *V*_*HC*<*SZ*_ mask, there are 26 networks where GIG-ICA shows significantly more negative t-stats in than STR, while in 5 networks out of 47, the t-stats are significantly more negative when estimated by STR. Fig. 5(C) shows that in strategy 3, for 34 out of 47 networks’ *V*_*HC*>*SZ*_ mask, the t-stats are significantly more positive when the networks are estimated by GIG-ICA. In contrast, superior parietal lobule, precuneus and left angular gyrus shows significantly higher (more positive) t-stats for STR. In case of the *V*_*HC*<*SZ*_ mask, there are 21 networks out of 47 where the t-stats are significantly more negative when estimated by GIGICA as opposed to 8 for STR.

Supplementary table S2 presents the exact number of discriminatory voxels found in networks estimated by each method following the three different strategies. It shows that that the highest number of significant voxels are identified in strategy 2, and the number of significant voxels in each network are different between GIG-ICA and STR. For strategy 3, the number of significant voxels is also different between the methods. This is because the process of identifying the significant voxels is independent between GIG-ICA and STR for these two strategies, with no overlap between the methods. Taken together, these results indicate that for the majority of the networks, GIG-ICA shows significantly greater group differences in the individual voxels with significant effect of diagnosis when compared to STR.

**Fig. 5:**
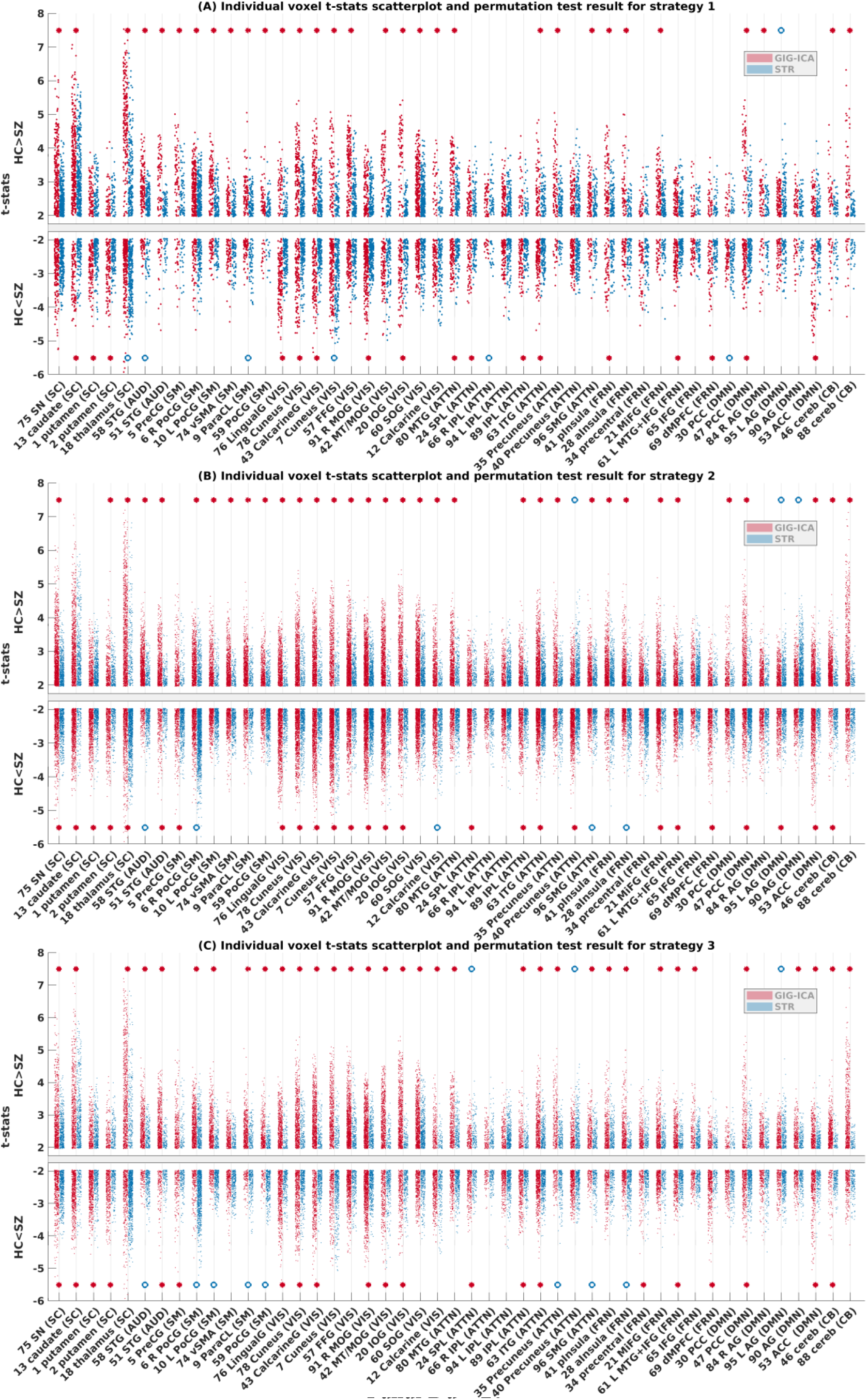
Individual voxel-based group difference and permutation test results, showing scatterplot of t-stats obtained from two-sample t-test between HC and SZ on each significant voxel within each network for strategy 1, 2 and 3 as outlined in section 2.2.4.1, in subfigures (A), (B) and (C) respectively. Permutation test was performed on voxels with positive t-stats and negative t-stats separately. For relevant networks, the red stars indicate that GIG-ICA shows significantly greater group difference than STR in the permutation test (*p* < 0.05), and the blue circles indicate that STR shows significantly greater group difference compared to GIG-ICA. The absence of such an indicator means that in that network there was no significant difference between GIG-ICA or STR estimated tstats. The full form name of each network can be found in supplementary table S1.

### 3.2 Classification results using the network maps

As mentioned above, we evaluated the classification ability using two datasets (FBIRN and COBRE). Fig. 6 shows the results with respect to FBIRN. For each method (STR or GIG-ICA) with each strategy (in feature selection), the accuracies in the testing data from 100 repeats of classification are shown in a column in Fig. 6(A) using a scatterplot, with a line, a darker box and a lighter box indicating the mean, 95% confident interval of mean and standard deviation respectively. The sensitivity and specificity are also shown in the same manner in Fig. 6(B) and 6(C), respectively. In addition, supplementary Fig. S4, S5 and S6 show the frequency of each voxel being used as a feature across 100 runs, grouped into 8 functional domains as estimated by the three strategies respectively. Fig. S7, S8 and S9 present the t-stats obtained from two-sample t-test between HCs and SZs in the training sets, averaged over 100 runs and the maximum in each functional domain in each voxel. Table S3 contains the average number of features in each functional domain over 100 runs of classification. For strategy 1, the average accuracy across 100 runs was 70.81±7.85% (sensitivity 0.69±0.1, specificity 0.74±0.11) for GIG-ICA and 64.97±7.82% (sensitivity 0.66±0.1, specificity 0.66±0.1) for STR. For strategy 2, the average accuracy was 76.45±8.9% (sensitivity 0.75±0.11, specificity 0.8±0.11) for GIG-ICA and 67.45±8.13% (sensitivity 0.65±0.11, specificity 0.71±0.11) for STR. For strategy 3, the accuracy was 72.65±7.96% (sensitivity 0.72±0.1, specificity 0.75±0.11) for GIG-ICA and 66.26±7.23% (sensitivity 0.65±0.09, specificity 0.68±0.1) for STR. To statistically evaluate the difference in the classification measures, two-sample t-tests were performed. The p-values obtained from the tests and noted in Fig. 6 are all significant at 0.05 level, which indicates that the GIG-ICA classification measures are all significantly higher than the STR measures. Hence it is evident that the average testing accuracy, sensitivity and specificity based on features estimated by GIG-ICA are higher than STR in all strategies.

**Fig. 6:**
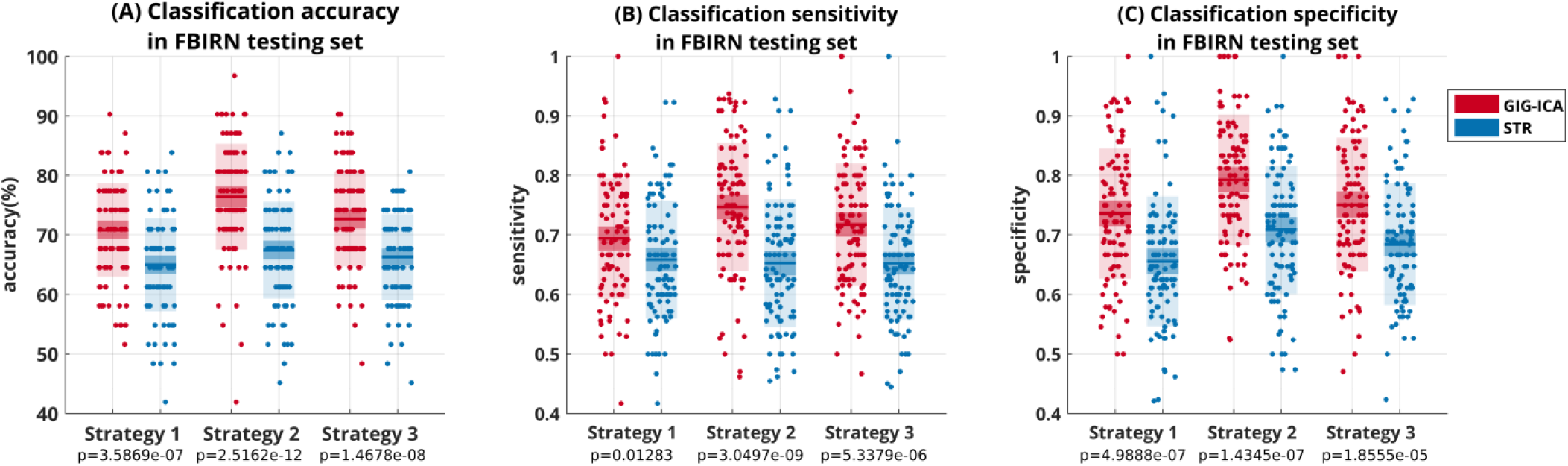
HC and SZ classification results obtained using networks estimated by GIG-ICA and STR methods from FBIRN data as features and SVM technique. Features were extracted from randomly selected training subjects using 3 different strategies and testing was performed on the remaining subjects. Classification with 10-fold cross-validation was repeated 100 times. Each dot represents accuracy in (A), sensitivity in (B) and specificity in (C), obtained in one of the 100 repeats. The horizontal line indicates mean result, darker box indicates 95% confidence interval of mean and lighter box indicates standard deviation of the results. The p-values obtained from two-sample t-tests between the GIG-ICA and STR measures for each strategy are mentioned below the x-axis.

The results above were obtained after including only the 47 resting-state networks in STR estimation. As noted in section 2.2.3, we also estimated STR by including all 100 group-level ICs. We then used the 47 subject-specific networks corresponding to the group-level resting state networks in the STR feature selection steps. Fig. 7 shows the classification results obtained using those. For strategy 1, the average accuracy across 100 runs was 62.55±8.26% (sensitivity 0.62±0.11, specificity 0.66±0.12) for STR. For strategy 2, the average accuracy was 63.68±7.67% (sensitivity 0.59±0.1, specificity 0.69±0.11) for STR. For strategy 3, the accuracy was 63.52±8.64% (sensitivity 0.61±0.1, specificity 0.68±0.1) for STR. It indicates that including the artifactual components does not improve the classification accuracy in STR. The focus of our work was to compare the efficacy of GIG-ICA and STR methods in estimating subject specific-networks using prior references. This experiment shows that STR does not perform better than GIG-ICA even if the ICs corresponding to artifacts, motion and other noise sources are included as the reference. On the contrary it fares worse compared to when only the resting-state networks are used as reference.

**Fig. 7:**
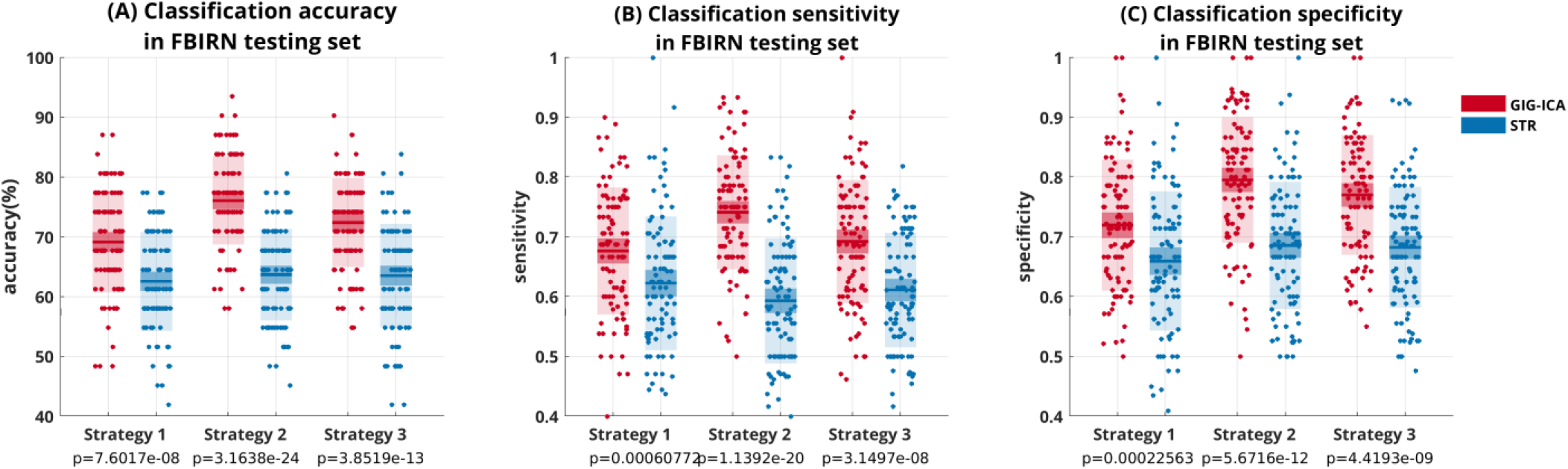
HC and SZ classification results obtained using networks estimated by GIG-ICA and STR methods from FBIRN data as features and SVM technique; notably the subject-specific STR networks were estimated while including the artifactual components as part of the reference. All the notations as well the GIG-ICA classification results are the same as in Fig. 6.

The superior classification performance of the GIG-ICA method generalizes well into the COBRE data, as demonstrated in Fig. 8. It is worth pointing out that the features were selected based on FBIRN data. Using the first scheme where the model was trained based on all FBIRN data, the predicted accuracy of the COBRE subjects using GIG-ICA and STR features were 78.66% and 75% respectively for strategy 1, 81.1% and 73.17% for strategy 2 and 79.88% and 73.17% for strategy 3. The sensitivity using GIG-ICA and STR features were 0.72 and 0.73 respectively for strategy 1, 0.73 and 0.72 for strategy 2 and 0.72 and 0.7 for strategy 3. The specificity using GIG-ICA and STR features were 0.85 and 0.77 respectively for strategy 1, 0.89 and 0.74 for strategy 2 and 0.88 and 0.77 for strategy 3. In the second scheme, SVM models were trained on randomly selected COBRE subjects using 10-fold cross-validation and then the rest were used in classification, which was repeated 10 times. For strategy 1, the average accuracy over the 10 runs was 83.13±11.04% (sensitivity 0.8±0.17, specificity 0.89±0.12) for GIG-ICA and 73.13±15.32% (sensitivity 0.73±0.22, specificity 0.78±0.18) for STR. For strategy 2, the average accuracy was 78.75±13.24% (sensitivity 0.77±0.15, specificity 0.84±0.17) for GIG-ICA and 70.63±11.04% (sensitivity 0.72±0.17, specificity 0.74±0.16) for STR. For strategy 3, the results were 76.25±7.1% (sensitivity 0.74±0.15, specificity 0.82±0.11) for GIG-ICA and 66.25±11.1% (sensitivity 0.66±0.18, specificity 0.7±0.13) for STR. The results indicate a clear trend toward higher classification performance using GIG-ICA estimated networks in predicting independent data.

**Fig. 8:**
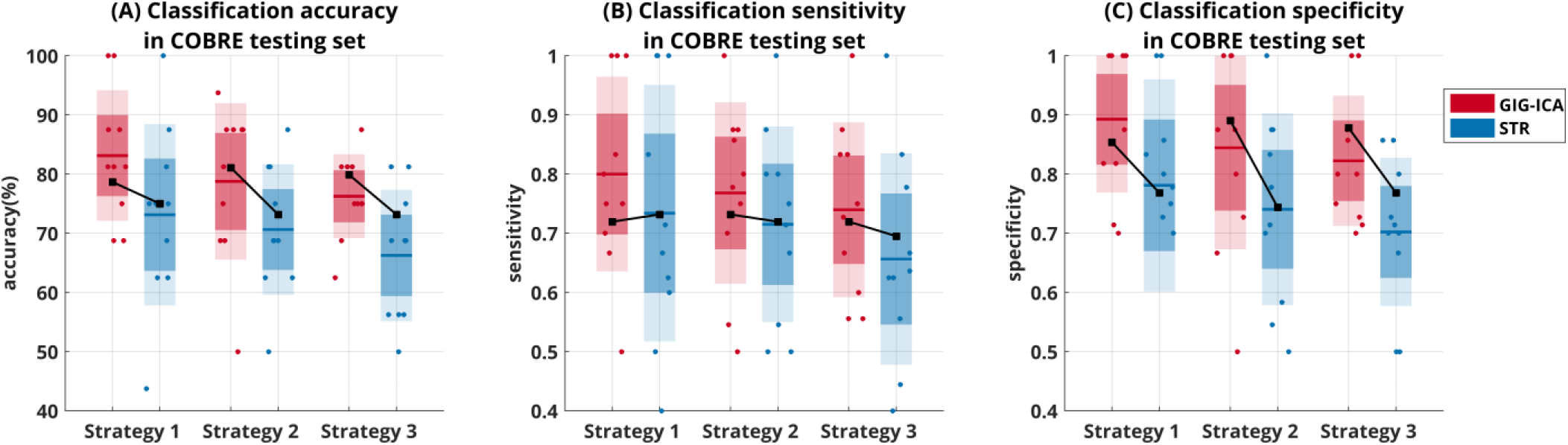
HC and SZ classification results of the independent dataset (COBRE). Features were extracted from subjects from the FBIRN dataset using 3 different strategies. The black lines in (A)-(C) indicate results from the first scheme where all FBIRN data were used for training and all COBRE data were used for testing the SVM model. In the second scheme, the classification was performed using a 10-fold cross-validation framework and was repeated 10 times. Each dot represents accuracy in (A), sensitivity in (B) and specificity in (C), obtained in one of the 10 repeats. The horizontal line indicates mean result, darker box indicates 95% confidence interval of mean and the lighter box indicates standard deviation of the results in the second scheme.

### 3.3 Comparison results of the functional network connectivity

In addition to the statistical analysis and classification on the subject-level spatially independent networks, we also examined the differences in FNC estimated by GIG-ICA and STR. Pairwise correlation between 47 networks’ TCs resulted in 47 × (47 − 1)/2=1081 connectivity values for each subject. Fig. 9(A) and 9(B) display the mean FNC across subjects estimated by GIG-ICA and STR, respectively. Notice that the GIG-ICA mean connectivity matrix appears to have higher contrast than the STR matrix. To establish this fact, a paired t-test was performed between each connectivity estimated using GIG-ICA and STR across all subjects. The t-stats obtained from the paired t-test on the connectivities are presented in Fig. 9(C). The positive t-stats indicate that connectivity strengths estimated using GIG-ICA have higher values compared to STR, and the negative t-stats indicate the opposite. It appears from the figure that the connectivity strengths estimated by GIG-ICA within the same domain (near the diagonal) are higher compared to STR.

**Fig. 9:**
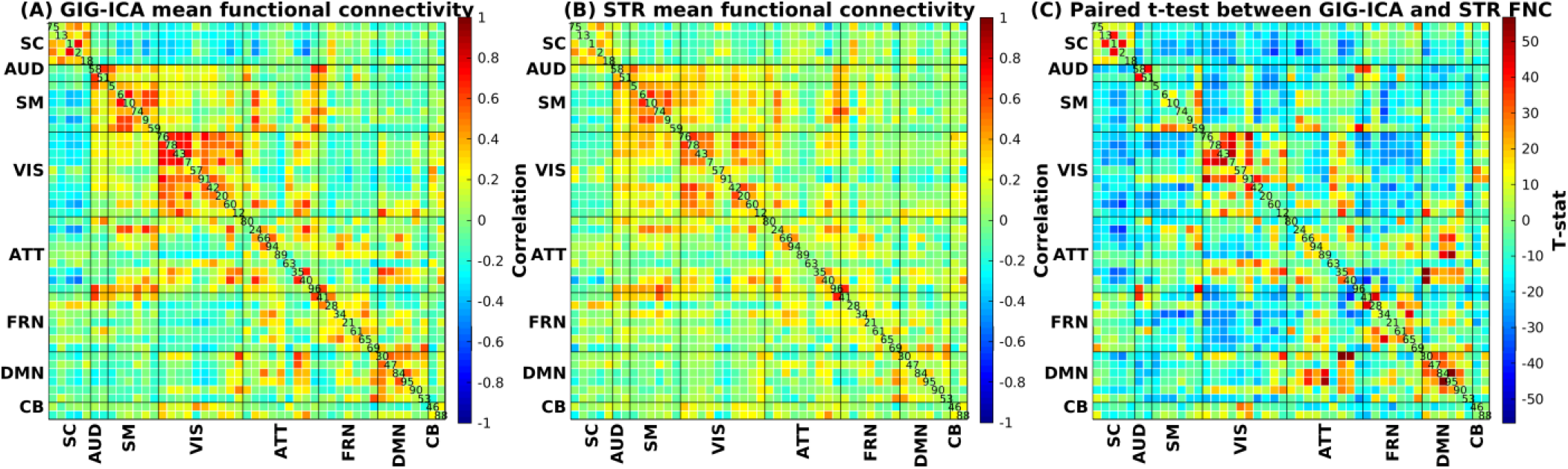
The mean FNC matrix across all subjects in FBIRN. (A) Mean FNC matrix estimated using GIG-ICA. (B) Mean FNC matrix estimated using STR. (C) Paired t-test result between the two methods based on the subject-specific FNC matrix.

We also compared the identified group difference between the two back-reconstruction methods. Fig. 10(A) and 9(B) show the t-stats obtained from two-sample t-tests performed on each connection estimated by GIG-ICA and STR respectively. Fig. 10(C) and 10(D) show the same t-stats thresholded at *p* < 0.05 after Bonferroni correction for multiple comparisons, for GIG-ICA and STR respectively. The thresholded FNC matrices indicate that there is considerable overlap between the group differences obtained from GIG-ICA and STR, but there are some important differences as well.

**Fig. 10:**
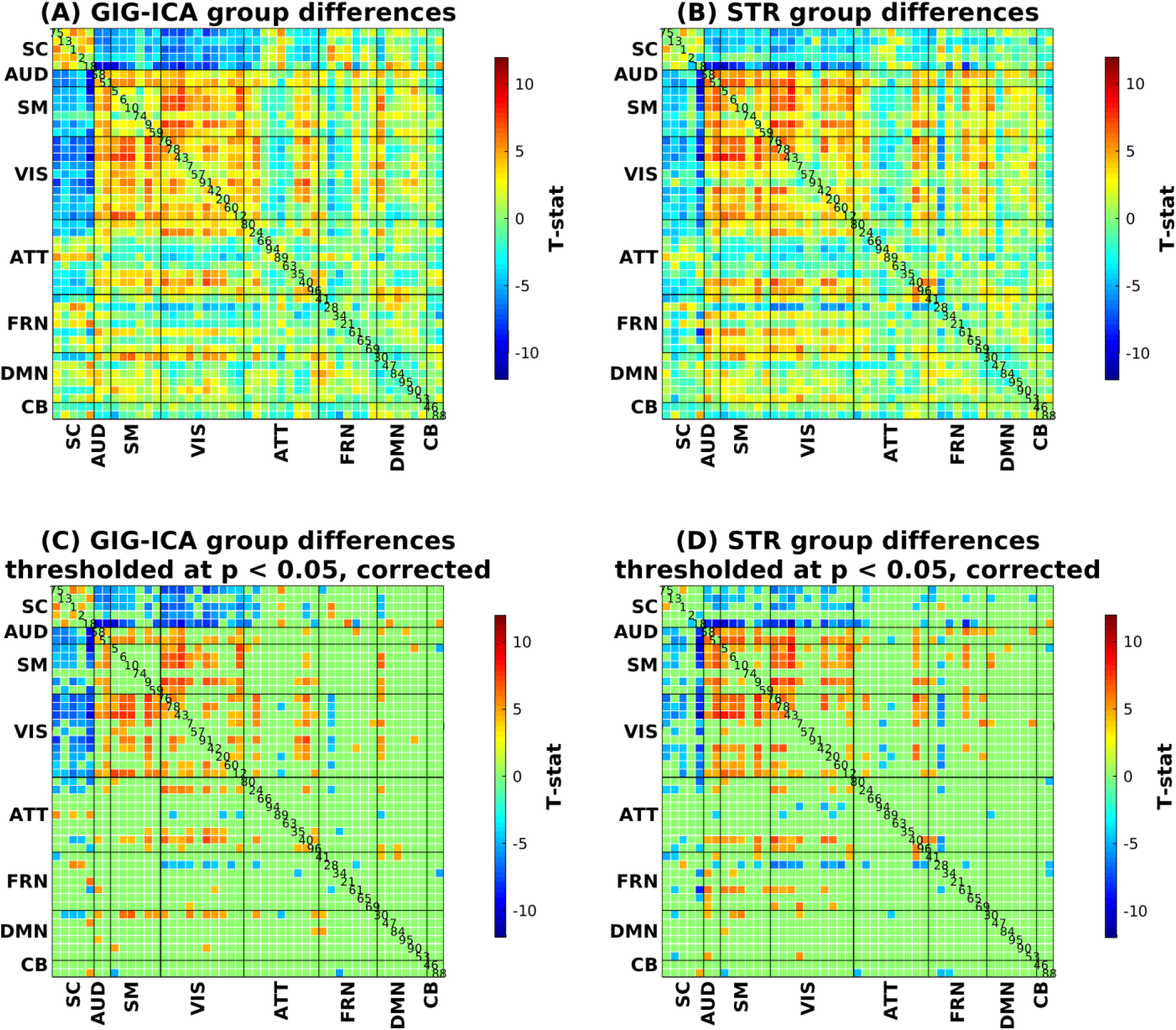
T-stats obtained from two-sample t-test between controls and patients in each element of the FNC matrix. (A) Group difference captured by GIG-ICA. (B) Group difference captured by STR. (C) Group difference captured by GIG-ICA, with t-stats thresholded at *p* < 0.05 after Bonferroni correction for multiple comparisons. (D) Group difference captured by STR, with t-stats thresholded at *p* < 0.05 after Bonferroni correction for multiple comparisons.

Since there were many connectivity pairs identified by both methods showing significant differences between the groups at 0.05 level of significance, a lower p-value threshold (10^−5^) was chosen to more clearly illustrate the differences. As shown in Fig. 11, both methods found unique alterations in schizophrenia. Fig. 11(A) shows the significant group differences identified by both methods. For both GIG-ICA and STR, SZs show stronger connectivity compared to HCs between thalamus (IC#18) and several sensory (auditory, visual and sensorimotor) networks. On the other hand, HCs show stronger connectivity than SZs between visual (left lingual, right cuneus and right calcarine gyrus) and sensorimotor (right postcentral, left precentral and left medial frontal gyrus) networks. Fig. 11(B) shows that at 10^−5^ level of confidence, connectivities differ significantly different between HC and SZ in 27 out of 1081 pairs for GIG-ICA but not STR. In fact, a look at supplementary Table S3 reveals that most of these FNC pairs were not significantly different between HC and SZ in the STR estimated result even at the 0.05 level. In contrast, 27 other FNC pairs out of 1081 were found to be significantly different at 10^−5^ level between HC and SZ in the STR result, as shown in Fig. 11(C), most of which were also significant in the GIG-ICA result at 10^−3^ level (see supplementary Table S4). These results suggest that GIG-ICA is more sensitive to group differences in the analysis of FNC than STR.

**Fig. 11:**
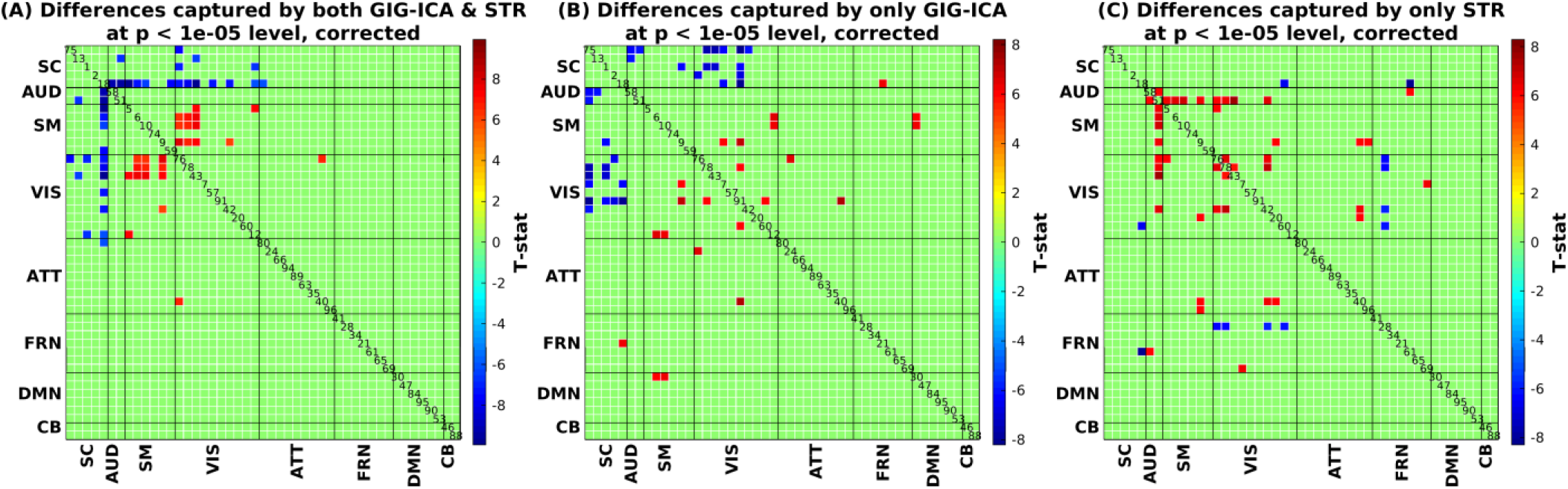
Significant group differences in FNC thresholded at *p* < 10^−5^ after Bonferroni correction for multiple comparisons. (A) Group differences captured by both GIG-ICA and STR. (B) Group differences captured by GIG-ICA only. (C) Group differences captured by STR only.

## 4 Discussion

In this study, we investigated the ability of different GICA back-reconstruction methods to identify biomarkers of schizophrenia. We compared GIG-ICA and STR methods in terms of the ability of the estimated subject-specific networks in differentiating HC and SZ groups by investigating the group difference in the network maps and the FNC as well as classifying patients from healthy controls using spatial networks. The STR method does not explicitly optimize independence of the subject-specific networks, and rather treats the component maps as fixed overlapping seeds (Joel et al., 2011). In contrast, the GIG-ICA method optimizes independence among the subject-specific components (Du et al., 2015b; Du and Fan, 2013). Recent studies have demonstrated the importance of incorporating individual variability in the estimation of functional networks and connectivity. Variation in the spatial arrangement of the functional brain regions across subjects directly translates into variability in the behavior and lifestyle of the individuals. In the absence of a measure which takes such variability into account in the fMRI data analysis, those can be wrongly interpreted as changes in functional connectivity (Bijsterbosch et al., 2018). Our findings in the current work reflect that the spatially constrained GIG-ICA method is more sensitive to group differences (even at the level of prediction of individual subjects from independent data), hence has more potential for biomarker identification.

Three different strategies were formulated to provide a fair comparison between the two methods. Strategy 1 allowed us to compare the group differences in the voxel z-scores and use the features in classification which were common in the two methods, with only the actual z-scores being different in the networks. Strategy 2 allowed for a feature selection and classification pipeline which was independent in the two methods. Strategy 3 considered the common positive activation (z-scores) in both methods; the subsequent step was independent. Note that only positive activation in voxels was considered and compared because voxels with positive activation represent more meaningful maps compared to the negative maps because the estimated components from ICA were converted to have positive kurtosis (Du et al., 2017a). The permutation tests on individual voxel group difference showed that for every strategy, t-stats computed based on GIG-ICA estimated networks in both *V*_*HC*>*SZ*_ and *V*_*HC*<*SZ*_ masks are significantly greater in majority of the 47 networks compared to STR.. Based on these results we conclude that more GIG-ICA-estimated networks, compared to STR, can indicate greater group difference.

Three different sets of features, i.e. the voxel z-scores for each component selected by the three slightly different strategies were used to classify HCs and SZs in the (primary) FBIRN data. The main observation from the classification results is that the average accuracy of classification based on features estimated by GIG-ICA method is higher than STR in all strategies, as indicated by the non-overlapping 95% confidence interval of mean in Fig. 6. Similar differences in the two methods are also evident in terms of the sensitivity and specificity metrics across all strategies. Strategy 2 employed a feature selection pipeline for the two methods which was independent from each other. As a result, the most discriminatory features were all included in the model training. Some of those features were lost in strategy 3, and more in strategy 1 because of the overlaps between two methods within the feature selection step. This explains why the difference in classification accuracy between the two methods was biggest in strategy 2.

Care was taken to avoid overfitting or underfitting the SVM classifier, and to generate models using an automatic feature selection pipeline which can show consistent result in new independent datasets. Our primary FBIRN dataset comprised of 163 HCs and 151 age and gender-matched SZs. The dimensionality of the fMRI data and the corresponding feature set was very high. In strategy 1, the average number of features used in model training was 8,140±445 across 100 repeats of classification. This number was the same for both GIG-ICA and STR, since we looked at common voxels in both methods in this strategy. In strategy 2, the average numbers of features were 52,551±2429 and 26,825±941 for GIG-ICA and STR respectively; and for strategy 3 those were 31,929±1317 and 19,466±699. Table S3 presents the average number of features in each functional domain over 100 runs of classification. Notice that the number of discriminating voxels between the controls and the patients is much higher in GIG-ICA estimated networks compared to STR, which is another advantage of this method. Although the dimensionality of the observations is very high, 314 observations is still a reasonably large sample size for a machine learning problem. SVM is not scale-invariant, hence every feature in the training set and the testing set were standardized separately (by setting mean=0 and standard deviation=1). In each repeat, model training and cross-validation was performed on 90% randomly selected subjects out of the whole dataset, and the performance of the trained model was reported based on prediction of the remaining subjects. Sensitivity and specificity are reported in addition to classification accuracy, as these measures take false positives and false negatives into account and thus are more desirable ways to communicate the performance of classification in identifying the target condition of interest (schizophrenia in this case). There are ways to formulate a more convincing testing procedure, such as using the trained models to predict classification performance on a different dataset containing HCs and SZs. In this paper, we evaluated this using the independent COBRE dataset, which also indicated higher classification accuracy and specificity when the features were estimated by GIG-ICA. In terms of sensitivity however, the two methods are very close, which is due to the nearly equal number of true positives detected.

The takeaway from the analysis of FNC is that at a higher significance level of 10^−5^, STR fails to capture some of the significant differences in connectivity between HCs and SZs. Connectivity between the functional networks is one of the emerging features of interest in diagnosis of mental illness. GIG-ICA generates visibly higher contrast in the mean FNC matrices and greater group difference in the two-sample t-test results. GIG-ICA also shows a higher number of group differences between subcortical networks and the other networks in which SZ patients have higher connectivity compared to HCs. GIG-ICA results also find more group differences between default mode network and other networks where HCs have higher connectivity compared to SZs. Note that in our study GIG-ICA is successful in identifying these results in schizophrenia using the average connectivity over the whole duration of the scan, whereas previous studies identified similar results in windowed dynamic FNC states (Damaraju et al., 2014a). These findings suggest the importance of using the higher-order statistical method to optimize the back-reconstruction at the single-subject level using spatially constrained ICA.

It should be noted that the STR subject network estimation could be improved by adding additional artifact remove steps and more aggressive motion correction (Griffanti et al., 2014), although artifact removal could improve the GIG-ICA network estimation as well. Another method for subject network estimation is template based rotation (TBR) which works by mapping data from new sessions into a priori templates (Schultz et al., 2014). TBR has shown larger group differences than dual regression and warrants a comparison with GIG-ICA in future. Future work may also focus on similar comparisons within the context of dynamic FNC analysis (Allen et al., 2014) and higher dimensional meta-states analysis of fMRI data (Miller et al., 2014). Another shortcoming in our study is that the FNC was not used as features for classification. We will investigate this issue in future since features from FNC can be used separately or in conjunction with the networks themselves for classification purpose (Silva et al., 2014).

While the same feature selection technique is used in the classification for both methods in the interest of fair comparison, the technique itself (i.e., one-sample t-test followed by two-sample t-test) used in this study is quite straightforward. This may have resulted in relatively lower overall classification accuracy across both methods and all strategies. While it is desirable for SVM (or any machine learning technique) to have as much data as possible to classify, more sensitive feature selection criteria (such as SVM recursive feature extraction (SVMRFE) (Guyon et al., 2002)) may result in better discriminatory features. In addition, SVM was the choice of classifier in this study due to its popularity in the field. More state-of-the-art feature selection and classification techniques (e.g., deep learning method) can be applied to improve classification performance.

This is an initial study on the efficacy of spatially-constrained ICA (e.g. GIG-ICA) methods in identifying fMRI-based biomarkers of schizophrenia. We found that this method is more sensitive to group differences and more powerful for the classification goal. Importantly, this approach enables ICA-based methods to scale to independent datasets by providing network correspondence between subjects while also optimizing for independence within the independent dataset. As GIG-ICA provides a promising approach for biomarker selection for schizophrenia, in future we will explore its ability using broader diseases such as bipolar disorder, Alzheimer’s disease and so on.

## Acknowledgements

This work was partially supported by National Institutes of Health grants 5P20RR021938/P20GM103472 and R01EB020407 and National Sciences Foundation grant 1539067 (to Calhoun VD), National Natural Science Foundation of China grant 61703253 and Natural Science Foundation of Shanxi grant 2016021077 (to Du YH).

